# growthcurves: A platform for human-in-the-loop analysis of biological growth curves

**DOI:** 10.64898/2026.05.23.727125

**Authors:** Samuel A. Bradley, Henry Webel, Stefano Donati, Carlos G. Acevedo-Rocha

**Affiliations:** The Novo Nordisk Foundation Biotechnology Research Institute for the Green Transition, Technical University of Denmark, DK-2800 Kongens Lyngby, Denmark

**Author notes:** To whom correspondence should be addressed: Stefano Donati, Carlos G. Acevedo-Rocha. These authors contributed equally to this work (Samuel A. Bradley, Henry Webel).

## Abstract

**Summary:** Biological growth curves are widely used but inconsistently analyzed due to fragmented workflows and limited quality control. We present *growthcurves*, a Python package for extracting growth parameters, and two open-source web applications, MicroGrowth and AutoGrowth, that combine automated fitting with interactive, human-in-the-loop inspection, selective refitting and traceable export for microplate reader and mini-bioreactor datasets in batch or turbidostat cultivation mode.

**Availability and Implementation:** growthcurves is implemented in Python and is freely available to non-commercial users at [https://github.com/biosustain/growthcurves.git] and through PyPI at [https://pypi.org/project/growthcurves/]. MicroGrowth and AutoGrowth are available at [https://biosustain.github.io/growthcurves_app/], and their source code is available at [https://github.com/biosustain/growthcurves_app.git]. Documentation, installation instructions, example datasets and tutorials are available at [https://growthcurves.readthedocs.io/en/latest/].

**Supplementary Information:** Supplementary information and Supplementary Methods are available online.

## Introduction

High-throughput time-series measurements of biological growth are widely used to quantify phenotypes across biological disciplines. Extracting meaningful parameters, such as intrinsic growth rate (μ), maximum specific growth rate (μ_max_), lag time (λ) and carrying capacity (K), requires careful preprocessing, model selection and quality control [1–6]. Numerous computational tools aid this process, including R packages [5,7,8], standalone programs [6], command-line tools [9] and web-based applications [10–12] (**Supp. Table S1**). However, despite differences in usability and flexibility, a common limitation is the separation of automated analysis from expert interpretation. This is particularly important as all approaches can produce fits that are mathematically valid but biologically implausible, particularly across diverse biological datasets, which vary in noise characteristics and growth dynamics. As a result, while recent tools facilitate iterative refitting with adjusted parameters [11], there is a general lack of integrated support and interactive diagnostics that enable rapid identification and correction of problematic fits across datasets. Consequently, analysis often becomes fragmented, with users switching between multiple platforms for data preprocessing, model fitting and visualization. In parallel, ambiguity in the interpretation of growth parameters, particularly distinctions between intrinsic and maximum growth rates, leads to inconsistent reporting [13,14]. Together, these challenges highlight the need for analysis frameworks that retain the scalability of automation while enabling transparent, user-driven validation within a single interface.

To address this, we developed growthcurves, a Python package, together with two open-source web applications, MicroGrowth and AutoGrowth. These tools implement a workflow in which automated fitting generates initial estimates, followed by interactive inspection and optional user-guided refinement through interactive visualizations. Users can then export the curated datasets, along with custom visualizations. This design supports the iterative, human-in-the-loop approach in a single platform that is needed for efficient and transparent generation of high-quality datasets by focusing and facilitating convenient quality control on problematic fits (**Supp. Fig. 1**).

### Python Package: *growthcurves*

The *growthcurves* Python package integrates established methods for growth curve analysis within a unified framework (**Supp. Methods**). It supports mechanistic parametric models for estimating intrinsic growth rate, lag time and carrying capacity, as well as phenomenological parametric and non-parametric approaches for estimating maximum specific growth rate (**Supp. Fig. S2, Supp. Table S2**). Non-parametric methods include spline fitting and sliding-window linear regression applied to log-transformed data. Two automatic options and a manual option are available for setting the spline smoothing factor, which strongly impacts μ_max_ estimates (**Supp. Fig. S3**,). All methods return a standardized data structure containing fitted parameters and derived statistics, enabling consistent downstream analysis (**Supp. Table S3**). Preprocessing functions, including blank subtraction, path length correction and outlier detection [15,16], are incorporated to improve robustness. Where growth phase boundaries are not directly returned by a method, it is estimated using threshold- or tangent-based approaches derived from instantaneous growth rates [11]. By consolidating these steps within a single framework, *growthcurves* reduces reliance on fragmented workflows spanning multiple tools.

### MicroGrowth Web Application

MicroGrowth is a convenient user-interface designed for plate-based assays (**Fig. 1a)**. Users upload time-series data alongside plate metadata and select *growthcurves* preprocessing and curve fitting options (**Supp. Fig. S4, S5**). Unlike previous GUIs, results are presented through comprehensive interactive visualizations where users can inspect global diagnostics, interrogate problematic fits and directly reanalyse wells with updated parameters, including refitting on highlighted subsections of the data (**Supp. Fig. S6**). Quality-controlled datasets can then be visualized as custom growth curve plots and comparative bar plots of calculated growth statistics (**Supp. Fig. S7**). By integrating automated analysis with interactive inspection in a single interface, MicroGrowth enables efficient yet quality-controlled analysis of plate experiments. Analysis settings and curated outputs can be exported as structured tables, supporting reproducible downstream analysis and transparent reporting [17].

**Figure 1.**
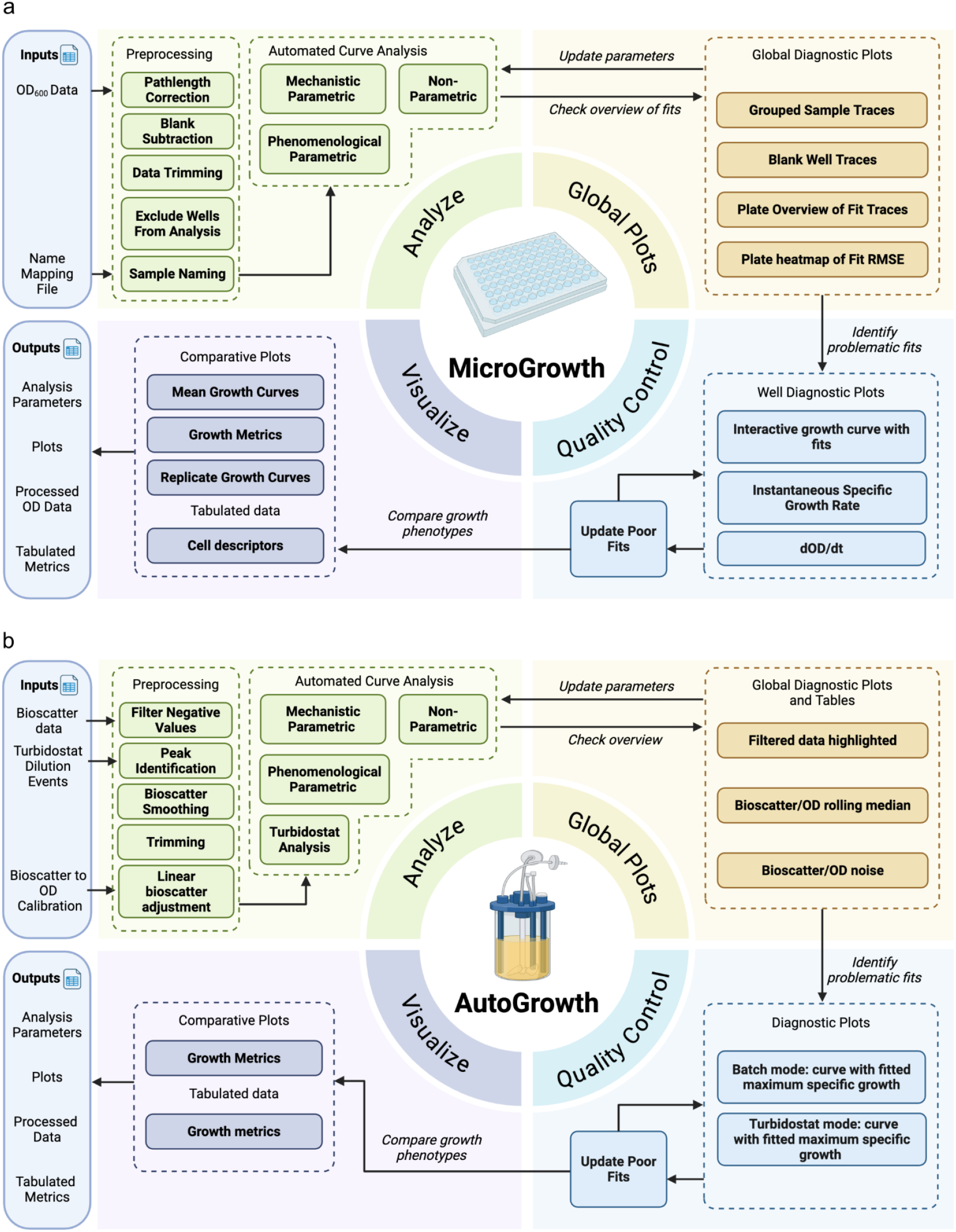
Growthcurves analysis functions and pipelines offered through the MicroGrowth and AutoGrowth front ends. **a**) Workflows for analyzing 96-well plate format data in the MicroGrowth application. **b**) Workflows for analyzing small-scale bioreactor data in the AutoGrowth application.

### AutoGrowth Web Application

AutoGrowth is tailored to data from automated mini-bioreactors such as the Chi.Bio [18] or Pioreactor (**Fig. 1b)**. In contrast to plate-based measurements, mini-bioreactor turbidity measurements are typically noisier and may vary across devices due to differences in sensor alignment. AutoGrowth thus incorporates smoothing, outlier detection and optional calibration against optical density measurements to enable quantitative comparison across experiments (**Supp. Fig. S8**). In addition, mini-bioreactors are often operated as turbidostats, producing repeated growth-dilution cycles rather than a single curve from a batch cultivation (**Supp. Fig. S9**). Turbidostat experiments can be used to maintain cells in metabolic steady-state for physiological studies or to automate adaptive laboratory evolution experiments. AutoGrowth supports thorough analysis of such data, through segmentation using recorded dilution metadata or automated detection of dilution events, enabling independent fitting of successive growth intervals (**Supp. Fig. S10**). As in MicroGrowth, results are visualized allowing users to validate and refine analyses prior to export

### Conclusion

Together, the *growthcurves* package and web applications address a key limitation in current growth-curve analysis workflows: the separation of automated computation from expert-driven intervention. Rather than assuming that automated fits are sufficient, the workflow makes points of uncertainty explicit and allows users to efficiently revise modelling decisions where needed. Integrating automated analysis and structured expert intervention in a single platform provides the flexibility for high quality analysis of complex and diverse biological datasets. This reduces reliance on *ad hoc* external analysis, making analysis decisions visible and traceable. As high-throughput phenotyping continues to expand, such human-in-the-loop approaches will help balance efficiency with biological interpretability, supporting more reliable quantitative characterization of growth.

### CRediT

S.A.B. and H.W. contributed equally to this work. S.A.B., S.D. and H.W. conceived the project. S.A.B. and H.W. developed the software, implemented the web applications and wrote the original draft. S.D. and C.G.A.-R. supervised the project and edited the manuscript. All authors reviewed and approved the final manuscript.

## Competing interests

The authors declare no competing interests.

## Funding

This work was supported by The Novo Nordisk Foundation [grant numbers NNF20CC0035580 and NNF24SA0100980] and Villum Fonden [grant number VIL78707].

## Acknowledgements

We thank Despoina Stavridou, Nils Emil Junge Marcussen, Frederick Hansson, Mehmet Mervan Cakar, Anna Maria Walerowicz, Catarina Rocha, Mariana Arango Saavedra, Baris Kara, Sergi Pujol Pinto and Felix Weichselbaum (Technical University of Denmark) for testing and feedback on the web applications and Magnus Ganer Jespersen for sharing an initial algorithm for turbidostat data analysis. Figures were created using BioRender.

## Notes

### Competing Interest Statement

The authors have declared no competing interest.

### Summary of Updates

Expanded supplementary methods. Added two new supplmentary figures (S2 and S3). reformatted the manuscript. Changed the manuscript title. Several small typo corrections.

## References

1. Tsoularis A, Wallace J. Analysis of logistic growth models. Math Biosci 2002;179:21–55.

2. Baranyi J, Roberts TA. A dynamic approach to predicting bacterial growth in food. Int J Food Microbiol 1994;23:277–94.

3. Huang L. A new mechanistic growth model for simultaneous determination of lag phase duration and exponential growth rate and a new Belehdrádek-type model for evaluating the effect of temperature on growth rate. Food Microbiol 2011;28:770–6.

4. Zwietering MH, Jongenburger I, Rombouts FM et al. Modeling of the bacterial growth curve. Appl Environ Microbiol 1990;56:1875–81.

5. Kahm M, Kahm M, Hasenbrink G et al. Grofit: Fitting biological growth curves. Derm Helv 2010, DOI: 10.1038/npre.2010.4508.

6. Hall BG, Acar H, Nandipati A et al. Growth rates made easy. Mol Biol Evol 2014;31:232–8.

7. Sprouffske K, Wagner A. Growthcurver: an R package for obtaining interpretable metrics from microbial growth curves. BMC Bioinformatics 2016;17:172.

8. Blazanin M. gcplyr: an R package for microbial growth curve data analysis. BMC Bioinformatics 2024;25:232.

9. Midani FS, Collins J, Britton RA. AMiGA: Software for automated analysis of microbial growth assays. mSystems 2021;6:e0050821.

10. Bukhman YV, DiPiazza NW, Piotrowski J et al. Modeling microbial growth curves with GCAT. Bioenergy Res 2015;8:1022–30.

11. Wirth NT, Funk J, Donati S et al. QurvE: user-friendly software for the analysis of biological growth and fluorescence data. Nat Protoc 2023;18:2401–3.

12. Reiter MA, Vorholt JA. Dashing Growth Curves: a web application for rapid and interactive analysis of microbial growth curves. BMC Bioinformatics 2024;25:67.

13. Ghenu A-H, Marrec L, Bank C. Challenges and pitfalls of inferring microbial growth rates from lab cultures. Front Ecol Evol 2024;11, DOI: 10.3389/fevo.2023.1313500.

14. Fernandez-Ricaud L, Kourtchenko O, Zackrisson M et al. PRECOG: a tool for automated extraction and visualization of fitness components in microbial growth phenomics. BMC Bioinformatics 2016;17:249.

15. Han S, Hu X, Huang H et al. ADBench: Anomaly Detection Benchmark. arXiv [csLG] 2022.

16. Chen S, Qian Z, Siu W et al. PyOD 2: A python library for outlier detection with LLM-powered model selection. Companion Proceedings of the ACM on Web Conference 2025. New York, NY, USA: ACM, 2025, 2807–10.

17. Meijer P, Howard N, Liang J et al. Provide proactive reproducible analysis transparency with every publication. R Soc Open Sci 2025;12:241936.

18. Steel H, Habgood R, Kelly CL et al. In situ characterisation and manipulation of biological systems with Chi.Bio. PLoS Biol 2020;18:e3000794.

