## Supplementary Information for "growthcurves: A platform for human-in-the-loop analysis of biological growth curves"

#### Abstract

**Summary:** Biological growth curves are widely used but inconsistently analyzed due to fragmented workflows and limited quality control. We present *growthcurves*, a Python package for extracting growth parameters, and two open-source web applications, MicroGrowth and AutoGrowth, that combine automated fitting with interactive, human-in-the-loop inspection, selective refitting and traceable export for microplate reader and mini-bioreactor datasets in batch or turbidostat cultivation mode. **Availability and Implementation:** growthcurves is implemented in Python and is freely available to non-commercial users at [https://github.com/biosustain/growthcurves.git] and through PyPI at [https://pypi.org/project/growthcurves/]. MicroGrowth and AutoGrowth are available at [https://biosustain.github.io/growthcurves_app/], and their source code is available at [https://github.com/biosustain/growthcurves_app.git]. Documentation, installation instructions, example datasets and tutorials are available at [https://growthcurves.readthedocs.io/en/latest/].

**Contact:**

### Supplementary Methods

#### *growthcurves*: Data preprocessing

The preprocessing subunit of the *growthcurves* package provides several preprocessing functions commonly used when analysing time-series growth data, including blank subtraction, OD correction for sample absorbances where the pathlength is not 1 cm and three outlier detection methods, namely interquartile range (IQR), the hampel method and the ECOD method [1,2].

#### *growthcurves*: Parametric and Non-Parametric Methods for Growth Curve Analysis

The *growthcurves* parametric module allows users to fit multiple mechanistic (Gompertz, Logistic, Richards, Baranyi) and phenomenological models (Gompertz, Logistic, Richards, Gompertz modified, Richards) via a single fit_parametric function with a model argument. The function employs scipy curve fit and returns a standardised output dictionary detailing the identity and fitted parameters of the chosen model.

The non_parametric module allows the user to apply a sliding window fitting of a linear model (sometimes known as the Easy Linear method) and spline curve fitting. As with the parametric fitting, both methods can be easily applied via a single fit_non_parametric function with a method argument. When fighting the spline curve, a cubic smoothing spline is fit to log-transformed OD via scipy.interpolate.make_smoothing_spline, then $\mu_{max}$ is taken as the maximum of the spline's first derivative. $\mu_{max}$ is sensitive to the λ (smoothing factor) (**Supp. Fig. 3**). growthcurves offers three methods for setting λ: 1) Manual, in which λ is fixed at a user-supplied value, 2) Heuristic (smooth="fast"), in which λ is set by a closed-form rule, $\lambda = c \cdot n \cdot \sigma^2$, where n is the number of timepoints and σ is Mean Absolute Deviation of the trimmed second differences of log-OD). The result is clipped to $[0, 0.8 \cdot n \cdot \mathrm{var}(\log OD)]$ and 3) Generalized Cross Validation (smooth="slow"), in which λ is chosen by generalized cross-validation.

#### *growthcurves*: Extracting phenotypic descriptors from fitted models

Phenotypic descriptors are extracted from the fitting function outputs by passing them to the extract_stats function, which returns a standardized dictionary containing all descriptors. **Supp. Table 3** details how each descriptor is calculated for each method. There are differences where some descriptors are fitted parameters, some are raw values and some are calculated via auxiliary functions. For example, in methods where the lag time is not returned by a fitted parameter, the lag time is calculated by either the threshold method, defined as the time at which the instantaneous specific growth rate increases above a user-specified fraction of the maximum specific growth rate, or the tangent method, defined as the time at which the tangent of the growth curve at the maximum specific growth rate intersects with the baseline OD.

#### AutoGrowth: Turbidostat Analysis

Optical density data from turbidostat cultures is first optionally pre-processed by filtering out downward-trending data points (negative OD changes) to reduce noise. The continuous culture data is then segmented into discrete growth intervals by identifying dilution events; these events are established either through imported bioreactor metadata or automated peak detection using scipy.signals.find_peaks with adjustable parameters for peak prominence and temporal distance (i.e. minimum peak height and temporal separation between peaks). Within each isolated growth segment, curve fitting is executed using the standard fitting functions from the *growthcurves* package, allowing for kinetic modeling via parametric and non-parametric approaches. Finally, by applying customizable phase boundary methods and thresholds, the software automatically delineates the exponential growth phases, calculates the maximum specific growth rate for each segment, and exports comprehensive statistical summaries and annotated growth plots.

#### *growthcurves*: Visualizing fits

The plot module provides convenient visualisation functions that take the standardised fit object, standardised descriptor object and any booleans for desired annotations as inputs and returns a Plotly plot with the data and requested annotations (e.g. calculated fit overlay, phase boundaries, point at $\mu_{max}$ and $OD_{max}$). The fit and descriptor objects can also be passed to the plot_derivative_metric function to plot instantaneous specific growth rate or the first derivative to the fit with similar annotations

***Supplementary Table 1. Comparison of available tools.*** *PM = parametric models, NP = non-parametric, QC = quality control, RMSE = root mean squared error, MSE = mean squared error, LOESS = locally estimated scatterplot smoothing, GUI = graphical user interface, RSS = residual sum of squares, AIC = Akaike information criterion.*

| Software | Code Base | Format | Curve Fitting Methods | Quality Control Options | Ref |
|---|---|---|---|---|---|
| grofit | CRAN<br><br>(archived) | R Package | **PM**: logistic, Gompertz, modified Gompertz, Richards.<br><br>**NP**: smooth spline fitting | **Readouts**: Graphical fit overlay; Fit QC (AIC, manual flagging of unreliable fits).<br><br>**Actions**: Scripted rerun. | [3] |
| GrowthRates | SourceForge | GUI | **PM**: none.<br><br>**NP**: sliding-window log-linear | **Readouts**: Graphical fit overlay of log-phase fit, Fit QC ($R^2$);<br><br>**Actions**: batch rerun | [4] |
| Growthcurver | GitHub | R Package | **PM**: logistic<br><br>**NP**: none | **Readouts**: Graphical Fit Overlay, Fit QC (residual standard error).<br><br>**Actions**: Scripted rerun | [5] |
| gcplyr | GitHub | R Package | **PM**: none.<br><br>**NP**: sliding-window log-linear and smooth spline fitting. | **Readouts**: Graphical Fit Overlay;<br><br>**Actions**: Scripted rerun | [6] |
| AMiGA | GitHub | CLI | **PM**: none.<br><br>**NP**: Gaussian Process Regression | **Readouts**: Graphical Fit Overlay; Fit QC (MSE, K-error)<br><br>**Actions**: CLI rerun. | [7] |
| GCAT | GitHub | GUI | **PM**: sigmoid curve (auto selection of | **Readouts**: Graphical Fit Overlay; Fit QC ($R^2$ and RSS) | [8] |

| | | | | | |
|---|---|---|---|---|---|
| | | | Richards, logistic or Gompertz)<br><br>**NP**: LOESS | **Actions**: batch rerun | |
| QurvE | GitHub | R Package + GUI | **PM**: logistic, Richards, Gompertz, extended Gompertz, Huang, Baranyi.<br><br>**NP**: sliding-window log-linear and smooth spline fitting. | **Readouts**: Graphical Fit Overlay; Fit QC ($R^2$, AIC and RMSE);.<br><br>**Actions**: batch rerun; Per-fit refit. | [9] |
| Dashing Growth Curves | GitHub | GUI | **PM**: logistic, Gompertz.<br><br>**NP**: sliding-window log-linear | **Readouts**: Graphical Fit Overlay; Fit quality ($R^2$ and RMSE).<br><br>**Actions**: Per-fit refit with option to manually select fit region | [10] |
| *growthcurves* + MicroGrowth/ PioGrowth | GitHub | Python Package + GUI | **PM**: logistic, Gompertz, modified Gompertz, Richards, Baranyi.<br><br>**NP**: smooth spline fitting and sliding-window log-linear | **Readouts**: Interactive Graphical Fit Overlay; Global plots for replicate consistency and fit quality (RMSE).<br><br>**Actions**: batch rerun; Per-fit refit with option to manually select exact datapoints; Manual adjustment, sample/well exclusion. | This work |

***Supplementary Table 2, Growth models: mechanistic and phenomenological models.*** *Parameters: $N_0$ initial population; $\mu$ growth rate; K carrying capacity; $\beta$ shape parameter; $h_0$ initial physiological state; $\lambda$ lag time; $\mu_{max}$ maximum growth rate; A asymptotic level; $\alpha$ secondary growth rate; t time $t_{shift}$ transition time; $\nu$ shape parameter. Models are formulated as in* [11]**.**

| Model | Equation | Key Parameters |
|---|---|---|
| **Mechanistic models** | | |
| Logistic (Mech) | $\frac{dN}{dt} = \mu\left(1 - \frac{N}{K}\right)N$ | $N_0, \mu, K$ |
| Gompertz (Mech) | $\frac{dN}{dt} = \mu \log\left(\frac{K}{N}\right)N$ | $N_0, \mu, K$ |
| Richards (Mech) | $\frac{dN}{dt} = \mu\left[1 - \left(\frac{N}{K}\right)^{\beta}\right]N$ | $N_0, \mu, \beta, K$ |
| Baranyi (Mech) | $\frac{dN}{dt} = \mu\frac{e^{\mu t}}{e^{h_0} - 1 + e^{\mu t}}\left(1 - \frac{N}{K}\right)N$ | $h_0, N_0, \mu, K$ |
| **Phenomenological models** | | |
| Logistic with lag (Phe) | $ln(N/N0) = \frac{A}{1 + \exp\left(\frac{4\mu_{\max}(\lambda - t)}{A} + 2\right)}$ | $\lambda, \mu_{\max}, A$ |
| Gompertz (Phe) | $ln(N/N0) = A\exp\left[-\exp\left(\frac{\mu_{\max}e}{A}(\lambda - t) + 1\right)\right]$ | $\lambda, \mu_{\max}, A$ |
| Modified Gompertz (Phe) | $ln(N/N0) = A\exp\left[-\exp\left(\frac{\mu_{\max}e}{A}(\lambda - t) + 1\right)\right] + A\exp\left(\alpha(t - t_{\text{shift}})\right)$ | $\lambda, \mu_{\max}, \alpha, t_{\text{shift}}, A$ |
| Richards (Phe) | $ln(N/N0) = A\left(1 + \nu\exp\left(1 + \nu + \frac{\mu_{\max}(1+\nu)^{1+1/\nu}}{A}(\lambda - t)\right)\right)^{-1/\nu}$ | $\lambda, \mu_{\max}, \nu, A$ |

***Supplementary Table 3. How key growth statistics are derived via each analysis method.*** *This table details how each growth metric is extracted from fitted model parameters or calculated from the raw data when each analysis method is applied. See Table 1 for parameters. Mech = mechanistic, phenom = phenomenological, np = non-parametric*

| Name | Lag Time | Intrinsic Growth Rate (μ) | Maximum Specific Growth Rate ($\mu_{max}$) | $OD_{max}$ | Fit Window |
|---|---|---|---|---|---|
| **Logistic (mech)** | threshold/tangent method | fitted μ | max dln(N)/dt of fitted model | fitted K | entire valid curve |
| **Gompertz (mech)** | threshold/tangent method | fitted μ | max dln(N)/dt of fitted model | fitted K | entire valid curve |
| **Richards (mech)** | threshold/tangent method | fitted μ | max dln(N)/dt of fitted model | fitted K | entire valid curve |
| **Baranyi (mech)** | fitted $h_0$ | fitted μ | max dln(N)/dt of fitted model | fitted K | entire valid curve |
| **Linear (np)** | threshold/tangent method | none | fitted gradient of linear line in sliding window | raw max OD | sliding window |
| **Spline (np)** | threshold/tangent method | none | max d spline(log OD) / dt | raw max OD | entire valid curve |
| **Logistic (phenom)** | fitted λ | none | fitted $\mu_{max}$ | max value of fitted curve within data range | entire valid curve |
| **Gompertz (phenom)** | fitted λ | none | fitted $\mu_{max}$ | max value of fitted curve within data range | entire valid curve |
| **Gompertz modified (phenom)** | fitted λ | none | fitted $\mu_{max}$ | max value of fitted curve within data range | entire valid curve |
| **Richards (phenom)** | fitted λ | none | fitted $\mu_{max}$ | max value of fitted curve within data range | entire valid curve |

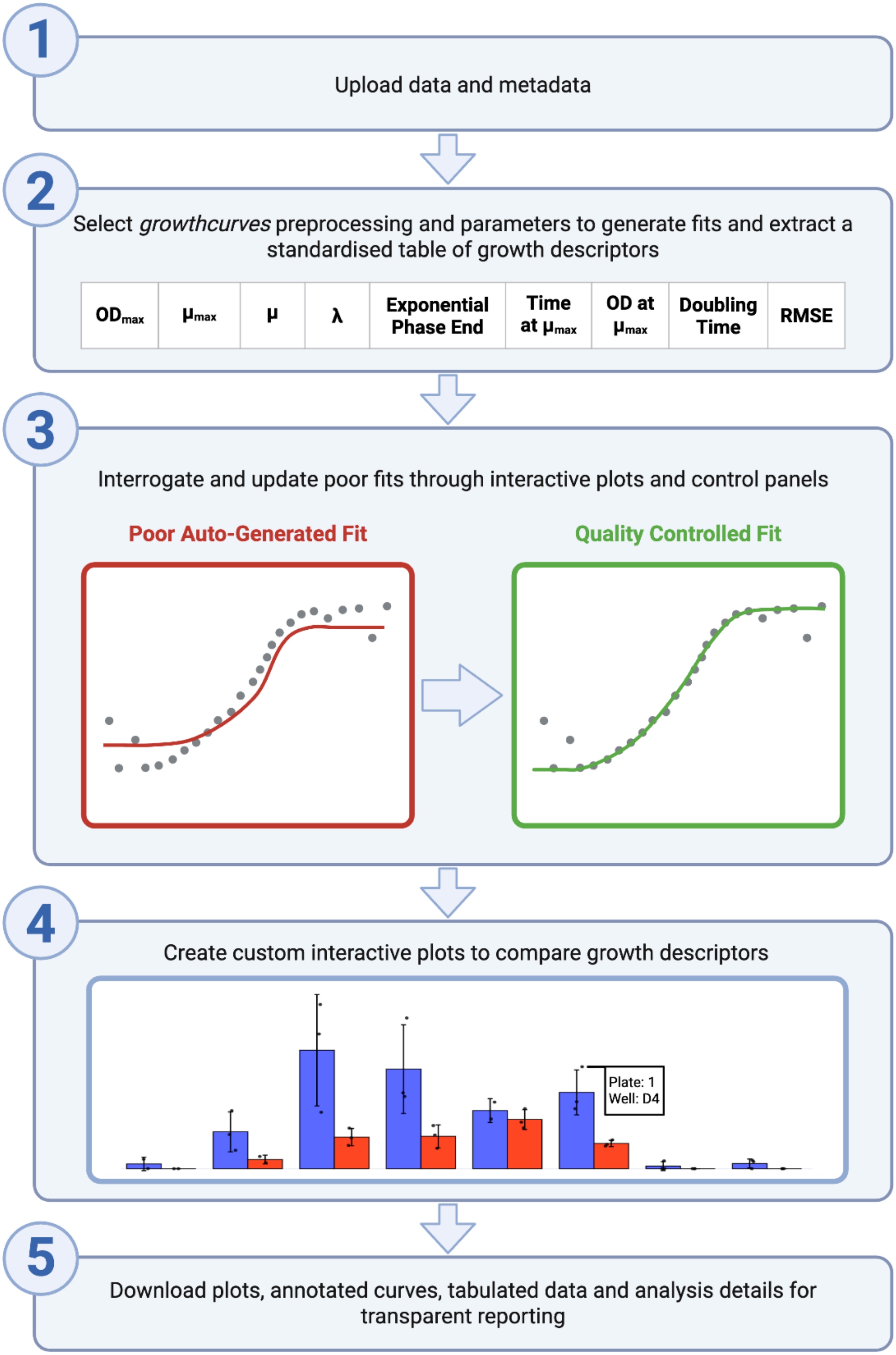
1
Upload data and metadata
2
Select *growthcurves* preprocessing and parameters to generate fits and extract a standardised table of growth descriptors
$OD_{max}$
$\mu_{max}$
$\mu$
$\lambda$
Exponential Phase End
Time at $\mu_{max}$
OD at $\mu_{max}$
Doubling Time
RMSE
3
Interrogate and update poor fits through interactive plots and control panels
Poor Auto-Generated Fit
Quality Controlled Fit
4
Create custom interactive plots to compare growth descriptors
Plate: 1
Well: D4
5
Download plots, annotated curves, tabulated data and analysis details for transparent reporting

***Supplementary Figure S1. Overview of the MicroGrowth and AutoGrowth workflow that support the human-in-the-loop growth curve analysis workflow.*** *(1) A user starts by uploading data and metadata files. (2) The user is guided to select the desired preprocessing and growthcurves analysis parameters for automatic curve fitting and calculation of a standardised set of growth descriptors. (3) The user can inspect individual growth curves to verify the fit quality and update the fitting parameters or directly update the descriptors if needed. (4) Once fit quality is confirmed, the user can generate comparative, interactive plots to gain biological understanding. (5) All plots, tabulated data and a history of analysis decision are available for download.*

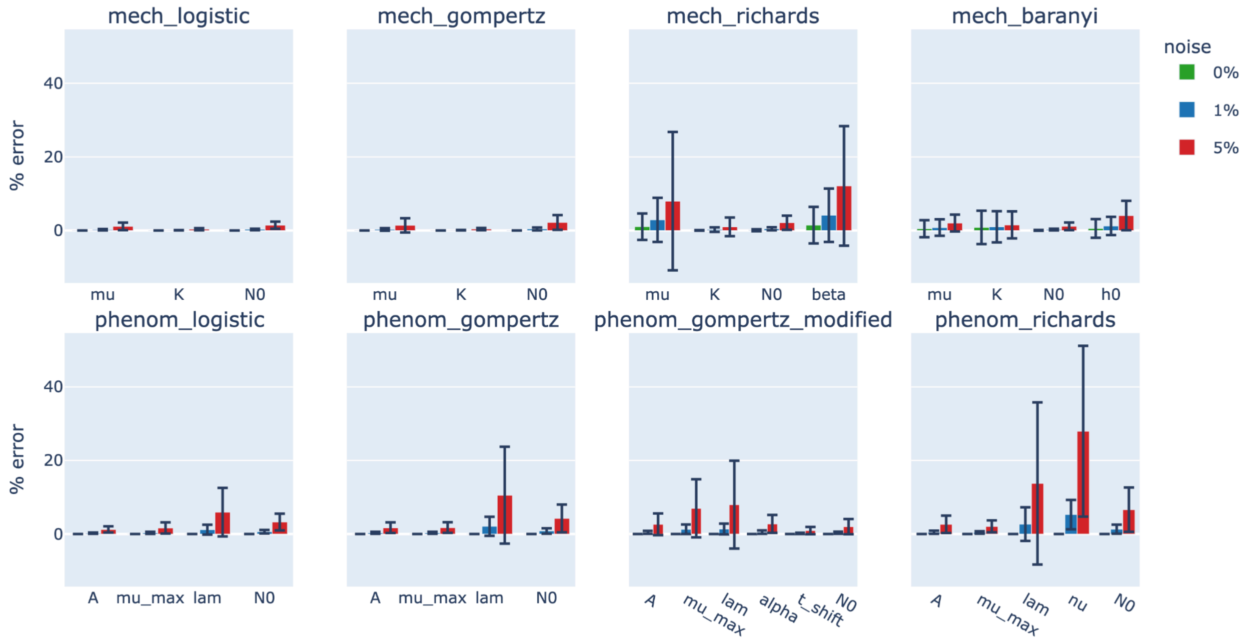


***Supplementary Figure S2. Parametric methods in growthcurves recover the parameters in synthetic growth curves.*** *Synthetic growth curves (24 h, 200 points) were generated from each model in growthcurves, corrupted with multiplicative Gaussian noise (σ = 0%, 1%, or 5%; green, blue, red), and fit with the same model. Bars show mean percentage error of each recovered parameter; error bars show ±1 SD across n = 50 replicates per condition.*

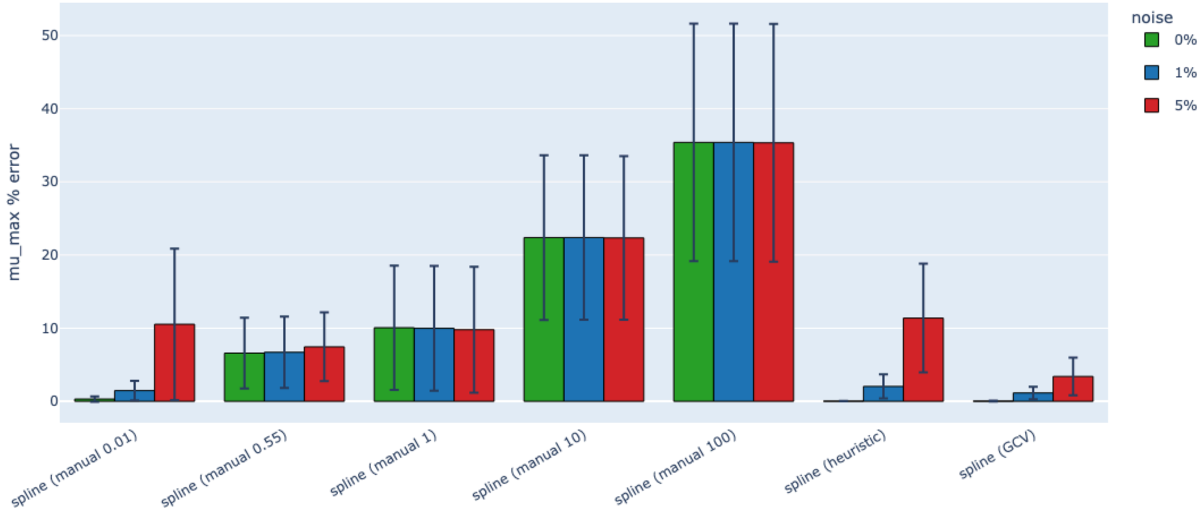


***Supplementary Figure S3. The accuracy and robustness of $\mu_{max}$ estimates are sensitive to the smoothing factor.*** *Fifty synthetic logistic growth curves were generated with multiplicative Gaussian noise and fit with the growthcurves spline method, using smoothing parameter (λ) fixed at manual values (0.01, 0.55, 1, 10, 100), by the automatic heuristic method, or by the automatic GCV method. Lower smoothing factors cause the spline to more closely follow data trends. Bars show the mean percentage error of $\mu_{max}$ estimates; error bars show ±1 SD.*

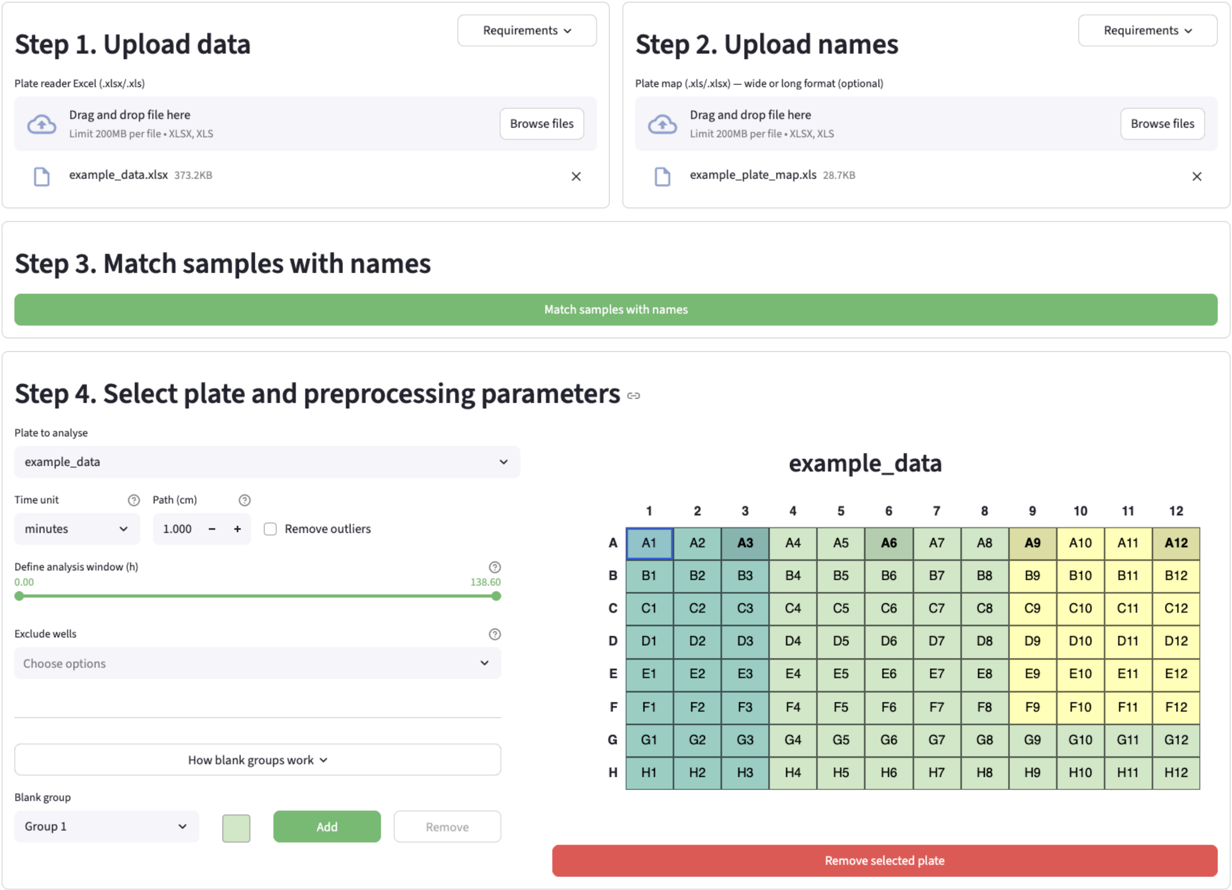


***Supplementary Figure S3. Schematic of upload options in MicroGrowth.*** *Users are guided through a stepwise process in which they upload tabulated growth data and a plate map naming file, select preprocessing options and define blanking groups (i.e. which blanks should be subtracted from which samples).*

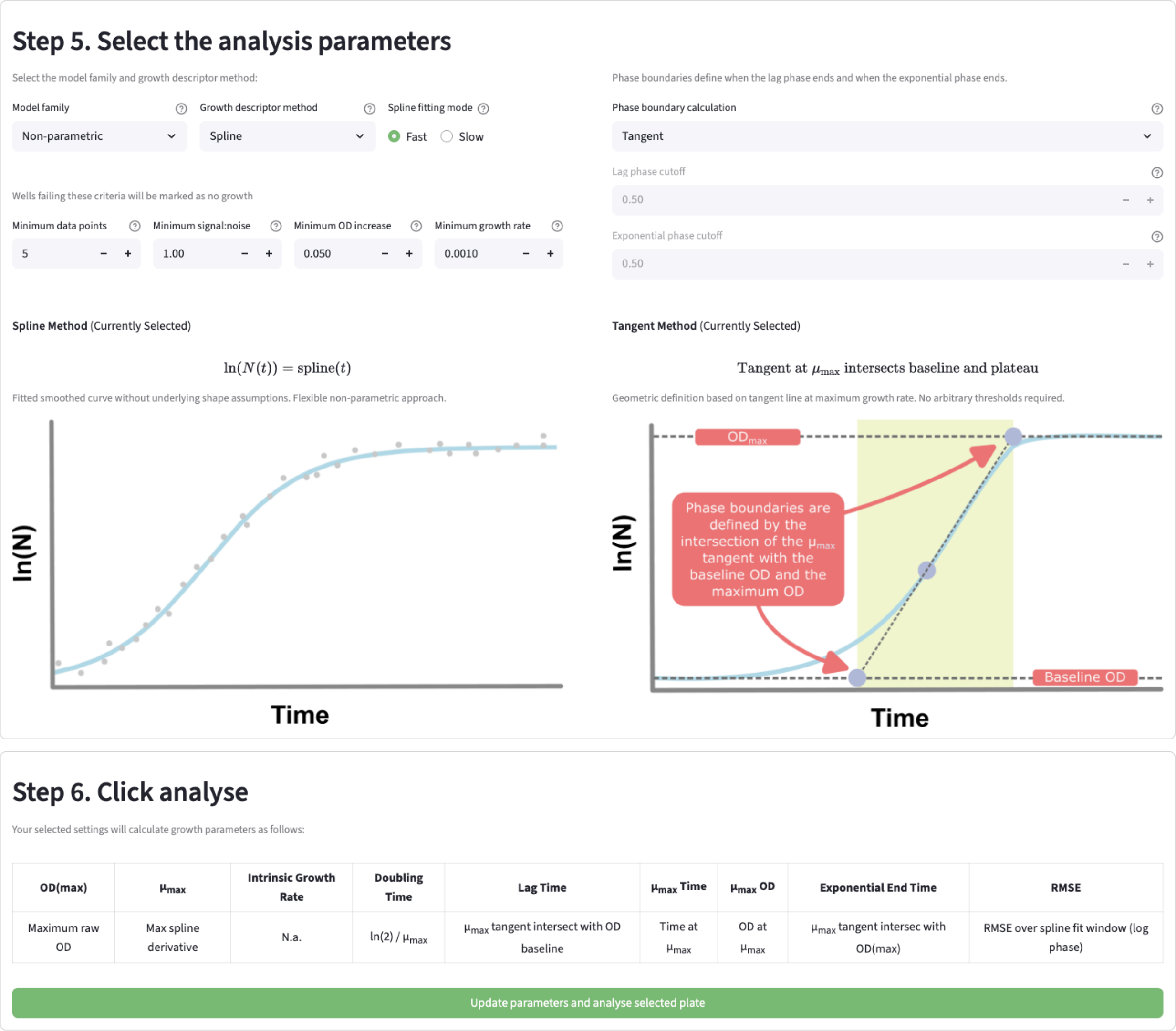


***Supplementary Figure 4. Schematic of analysis options in MicroGrowth.*** *Users select the analysis parameters for growth curve analysis, including which parametric or non-parametric model to use, thresholds for identifying a well as having 'No Growth' and how the lag time should be calculated if the lag time is not a fitted parameter. A tabulated summary of how each parameter is calculated for the selected parameter set is displayed.*

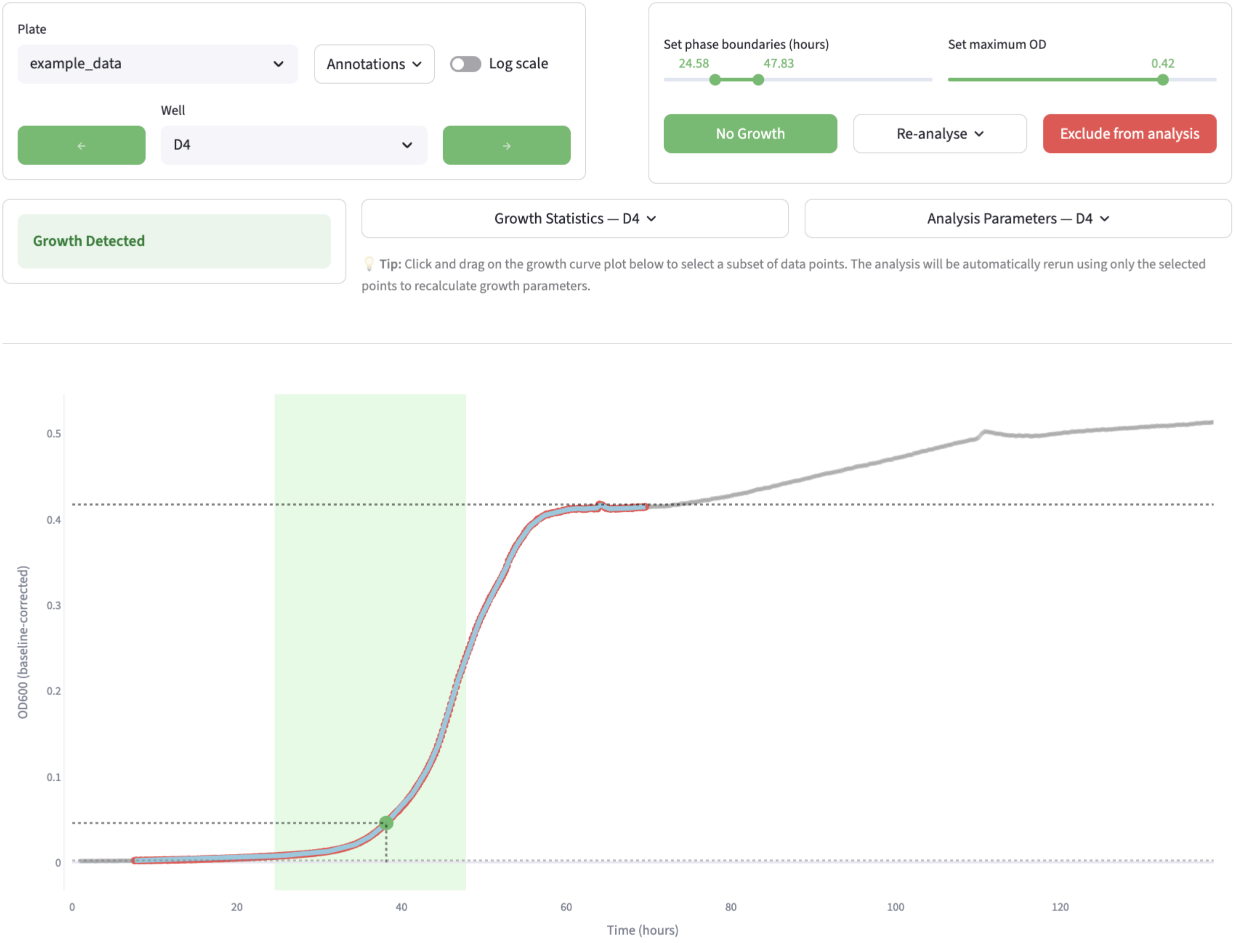


***Supplementary Figure 5. Schematic of diagnostic fit plots in MicroGrowth.*** *Users navigate through fitted growth curves and can visually assess and update the fit quality through interactive graphs. Individual wells can be quickly reanalyzed with updated parameters and specific subsections of the data can be reanalyzed by selecting points in the interactive graph. This is useful for avoiding noisy stationary phase data, which can otherwise impact fit quality.*

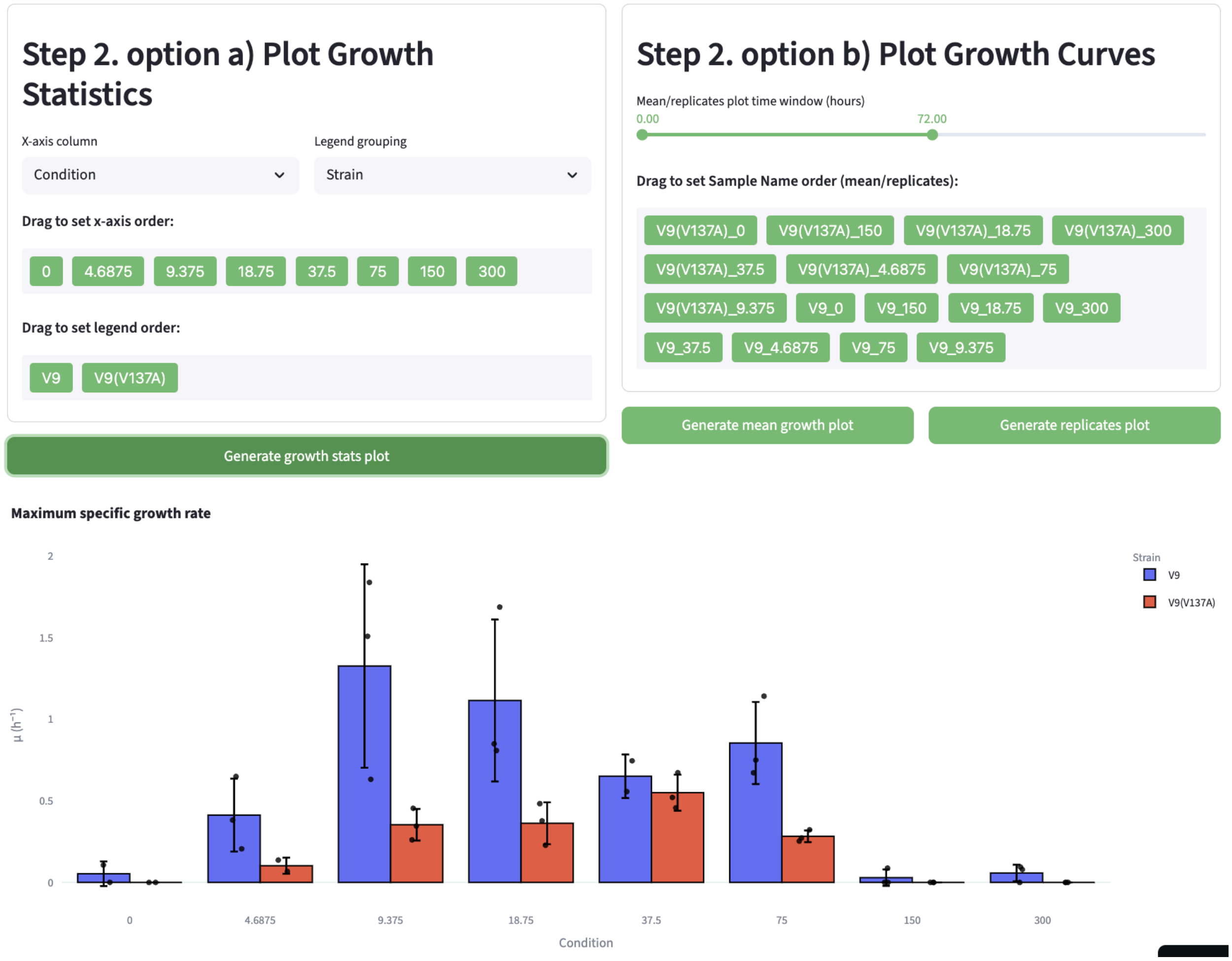


***Supplementary Figure 6. Schematic of visualization of calculated growth descriptors in MicroGrowth.*** *Users are guided through a stepwise process in which they first select the wells they wish to include (Step 1, not shown). Users can then generate grouped bar plots to compare calculated growth descriptors as well as plots of growth curves vs time (either as the mean trace of replicates or as individual replicates). Bars represent mean values ±1 SD.*

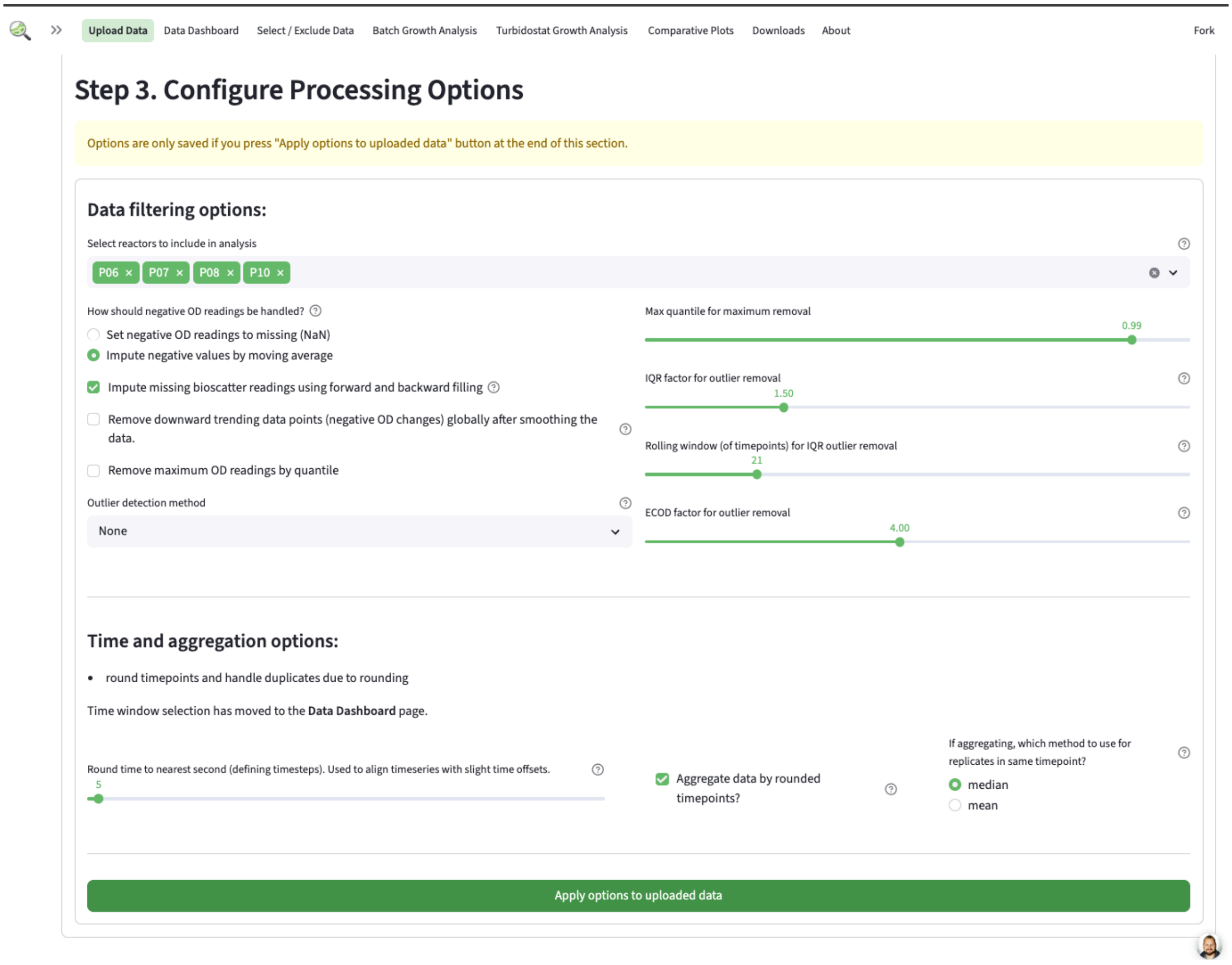


***Supplementary Figure 7. Data filtering options for noisy turbidity measurements from optical devices in AutoGrowth.*** Different options can be selected and configured. Per default negative measurements are imputed using the moving average. Additionally missing values can be replaced by forward and then backward filling. Removal of downward trending data points is especially useful for turbidostat experiments to get clean growth curves for each time interval. To align the measurements across reactors with slight offsets, the time is rounded to the next 5 seconds, which can be adjusted. If the rounding leads to duplicates, an aggregation can be performed.

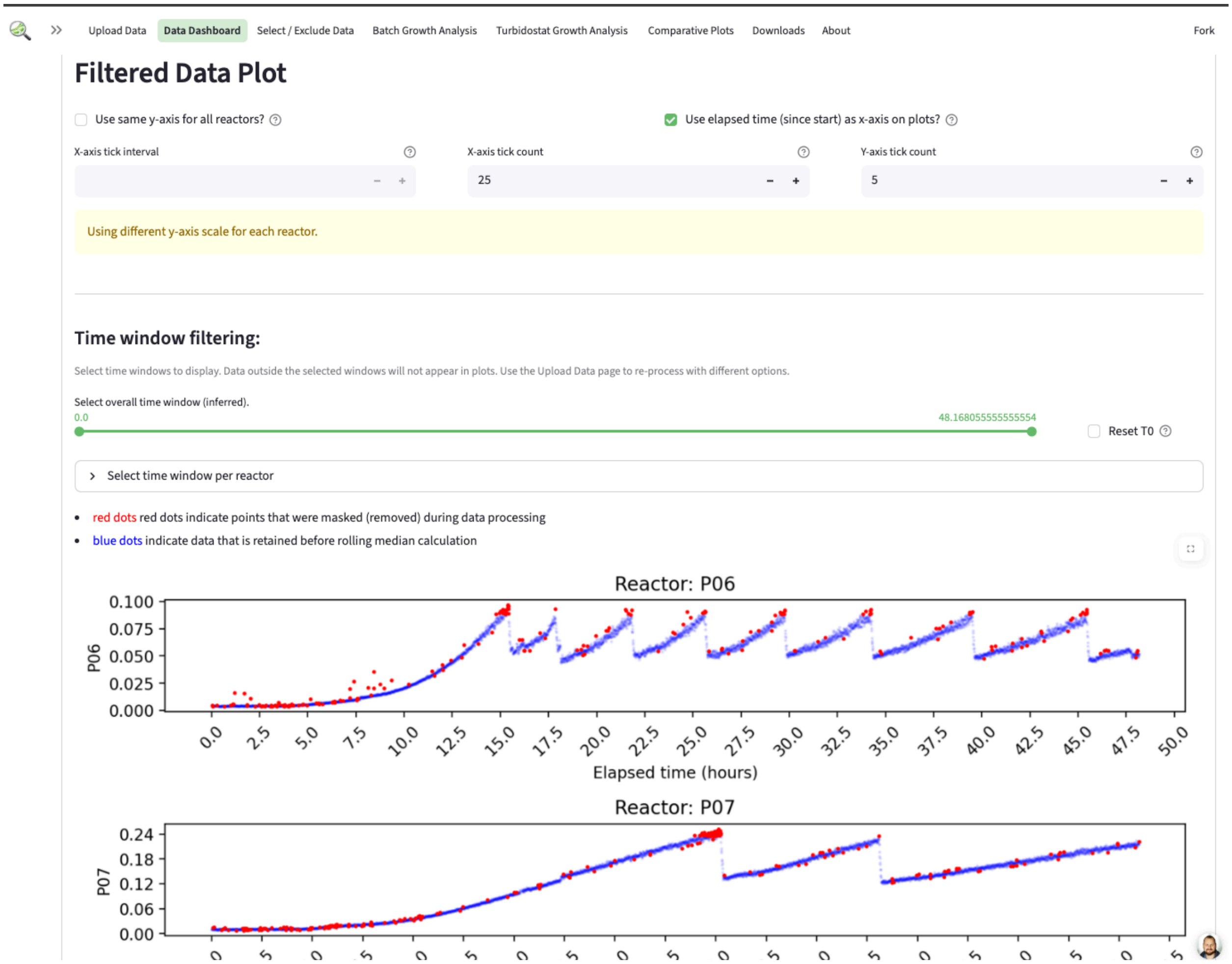


***Supplementary Figure 8. Overview of data after filtering in AutoGrowth.*** *After filtering the data is presented in a dashboard where measurements can be trimmed, per default, without changing the absolute zero timepoint of the start of the experiment. Filtered data points are highlighted in red.*

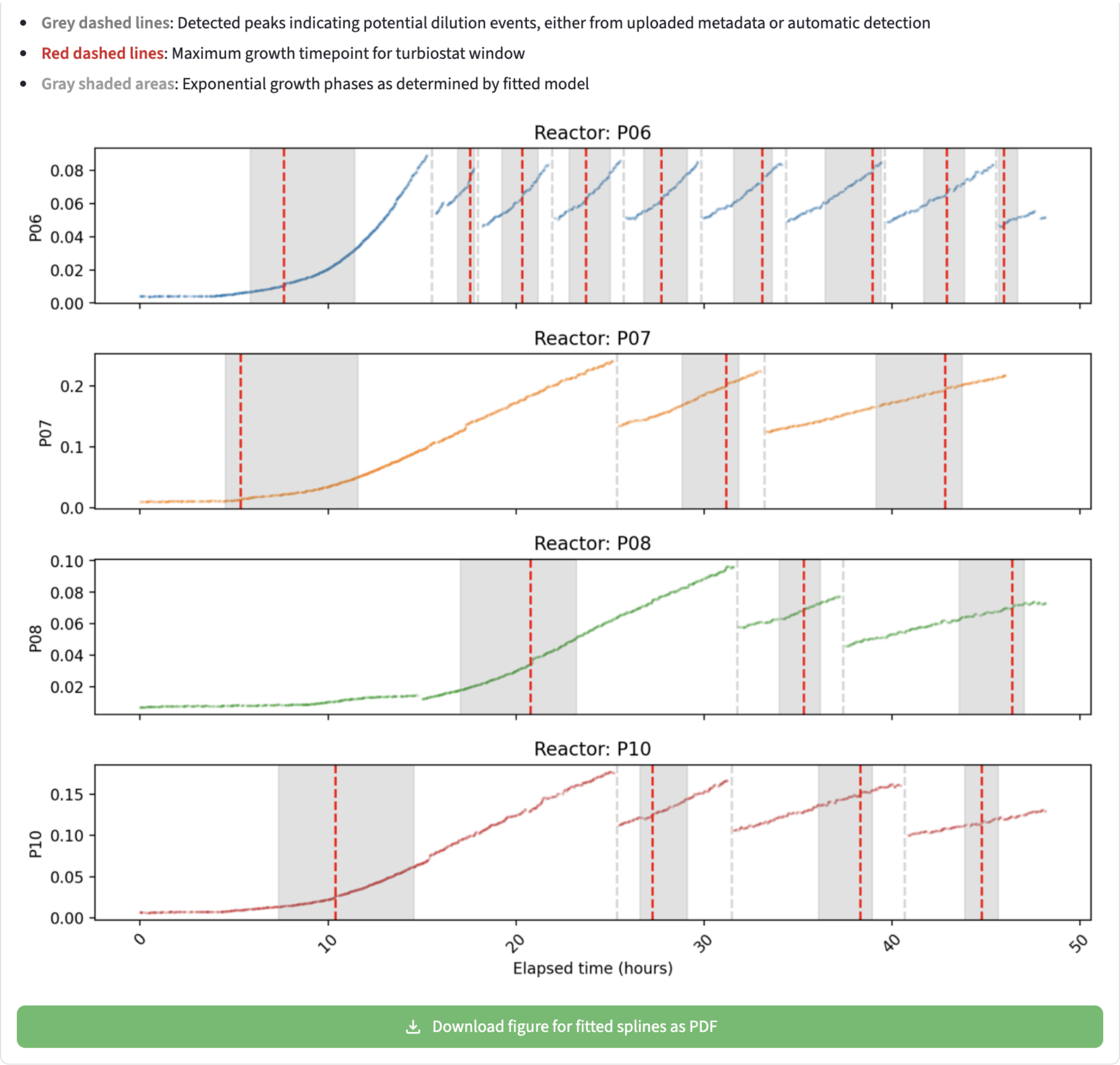


***Supplementary Figure 9. Exponential growth phases per interval in AutoGrowth.*** *The user can upload dilution events for turbidostat mode experiments or use automated peak detection to determine the intervals. The app then visualizes the maximum growth per interval. The statistics are provided as a table below the figure in which each turbidostat interval in each reactor is a single row.*

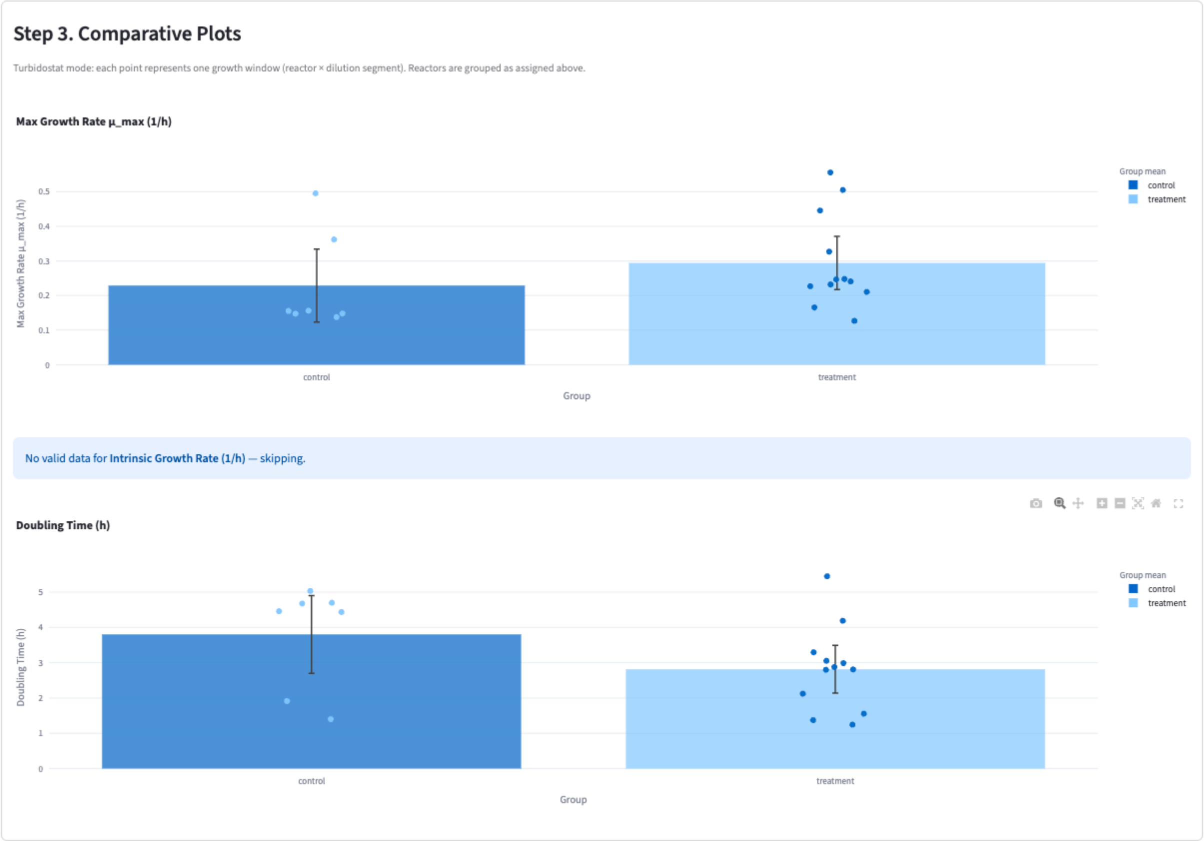


***Supplementary Figure 10. Comparative plots of calculated statistics per group in AutoGrowth.*** *The user can group samples in the app and then visualize all the calculated statistics in a separate page. Bars represent mean values ±1 SD. Each turbidostat interval is a single dot in the comparison plot.*

### References


1. Han S, Hu X, Huang H *et al.* ADBench: Anomaly Detection Benchmark. *arXiv [csLG]* 2022.

2. Chen S, Qian Z, Siu W *et al.* PyOD 2: A python library for outlier detection with LLM-powered model selection. *Companion Proceedings of the ACM on Web Conference 2025*. New York, NY, USA: ACM, 2025, 2807–10.

3. Kahm M, Hasenbrink G, Lichtenberg-Fraté H *et al.* grofit: Fitting Biological Growth Curves withR. *J Stat Softw* 2010;**33**, DOI: 10.18637/jss.v033.i07.

4. Hall BG, Acar H, Nandipati A *et al.* Growth rates made easy. *Mol Biol Evol* 2014;**31**:232–8.

5. Sprouffske K, Wagner A. Growthcurver: an R package for obtaining interpretable metrics from microbial growth curves. *BMC Bioinformatics* 2016;**17**:172.

6. Blazanin M. gcplyr: an R package for microbial growth curve data analysis. *BMC Bioinformatics* 2024;**25**:232.

7. Midani FS, Collins J, Britton RA. AMiGA: Software for automated analysis of microbial growth assays. *mSystems* 2021;**6**:e0050821.

8. Bukhman YV, DiPiazza NW, Piotrowski J *et al.* Modeling microbial growth curves with GCAT. *Bioenergy Res* 2015;**8**:1022–30.

9. Wirth NT, Funk J, Donati S *et al.* QurvE: user-friendly software for the analysis of biological growth and fluorescence data. *Nat Protoc* 2023;**18**:2401–3.

10. Reiter MA, Vorholt JA. Dashing Growth Curves: a web application for rapid and interactive analysis of microbial growth curves. *BMC Bioinformatics* 2024;**25**:67.

11. Ghenu A-H, Marrec L, Bank C. Challenges and pitfalls of inferring microbial growth rates from lab cultures. *Front Ecol Evol* 2024;**11**, DOI: 10.3389/fevo.2023.1313500.
